# Structured Reviews for Data and Knowledge Driven Research

**DOI:** 10.1101/729475

**Authors:** Núria Queralt-Rosinach, Gregory S. Stupp, Tong Shu Li, Michael Mayers, Maureen E. Hoatlin, Matthew Might, Benjamin M. Good, Andrew I. Su

## Abstract

**Motivation:** Hypothesis generation is a critical step in research and a cornerstone in the rare disease field. Research is most efficient when those hypotheses are based on the entirety of knowledge known to date. Systematic review articles are commonly used in biomedicine to summarize existing knowledge and contextualize experimental data. But the information contained within review articles is typically only expressed as free-text, which is difficult to use computationally. Researchers struggle to navigate, collect and remix prior knowledge as it is scattered in several silos without seamless integration and access. This lack of a structured information framework hinders research by both experimental and computational scientists.

**Results:** To better organize knowledge and data, we built a structured review article that is specifically focused on NGLY1 Deficiency, an ultra-rare genetic disease first reported in 2012. We represented this structured review as a knowledge graph, and then stored this knowledge graph in a Neo4j database to simplify dissemination, querying, and visualization of the network. Relative to free-text, this structured review better promotes the principles of findability, accessibility, interoperability, and reusability (FAIR). In collaboration with domain experts in NGLY1 Deficiency, we demonstrate how this resource can improve the efficiency and comprehensiveness of hypothesis generation. We also developed a read-write interface that allows domain experts to contribute FAIR structured knowledge to this community resource. In contrast to traditional free-text review articles, this structured review exists as a living knowledge graph that is curated by humans and accessible to computational analyses. Finally, we have generalized this workflow into modular and repurposable components that can be applied to other domain areas. This NGLY1 Deficiency-focused network is publicly available at http://ngly1graph.org/.

**Availability and implementation:** Source code and network data files are at: https://github.com/SuLab/ngly1-graph and https://github.com/SuLab/bioknowledge-reviewer.

## Introduction

Science progresses via an iterative loop between hypothesis generation, experimentation, and interpretation. Interpretation relies on putting new data in context with existing relevant knowledge. Researchers typically need to access the relevant knowledge to their research question and hypothesis. One method we have for accessing existing relevant knowledge are review articles, as they organize knowledge around a topic. These systematic review articles are common in biomedicine, but the content is typically expressed only as free-text, which is not easily queryable and computable. Consequently, the community does not benefit from the full value of review articles.

As a result, the informatics community is very often faced with the challenge of integrating data across many knowledge resources (**Figure 1**). Most of these knowledge resources organize information for a relatively shallow scope of information types, and across a wide breadth of biological areas. For example, BioGRID focuses on integrating data on physical and genetic interactions (1), and the Gene Ontology Consortium annotates functions of gene products (2). In addition, information aggregation platforms like Monarch (3), the EBI RDF platform (4), and Open PHACTS (5) integrate data for several information types into a single resource. While these meta-databases are valuable, their objective is typically different from the goals of a disease-specific review article, which usually focus on a richly heterogeneous network of knowledge in a relatively limited scope of biology.

**Figure 1.**
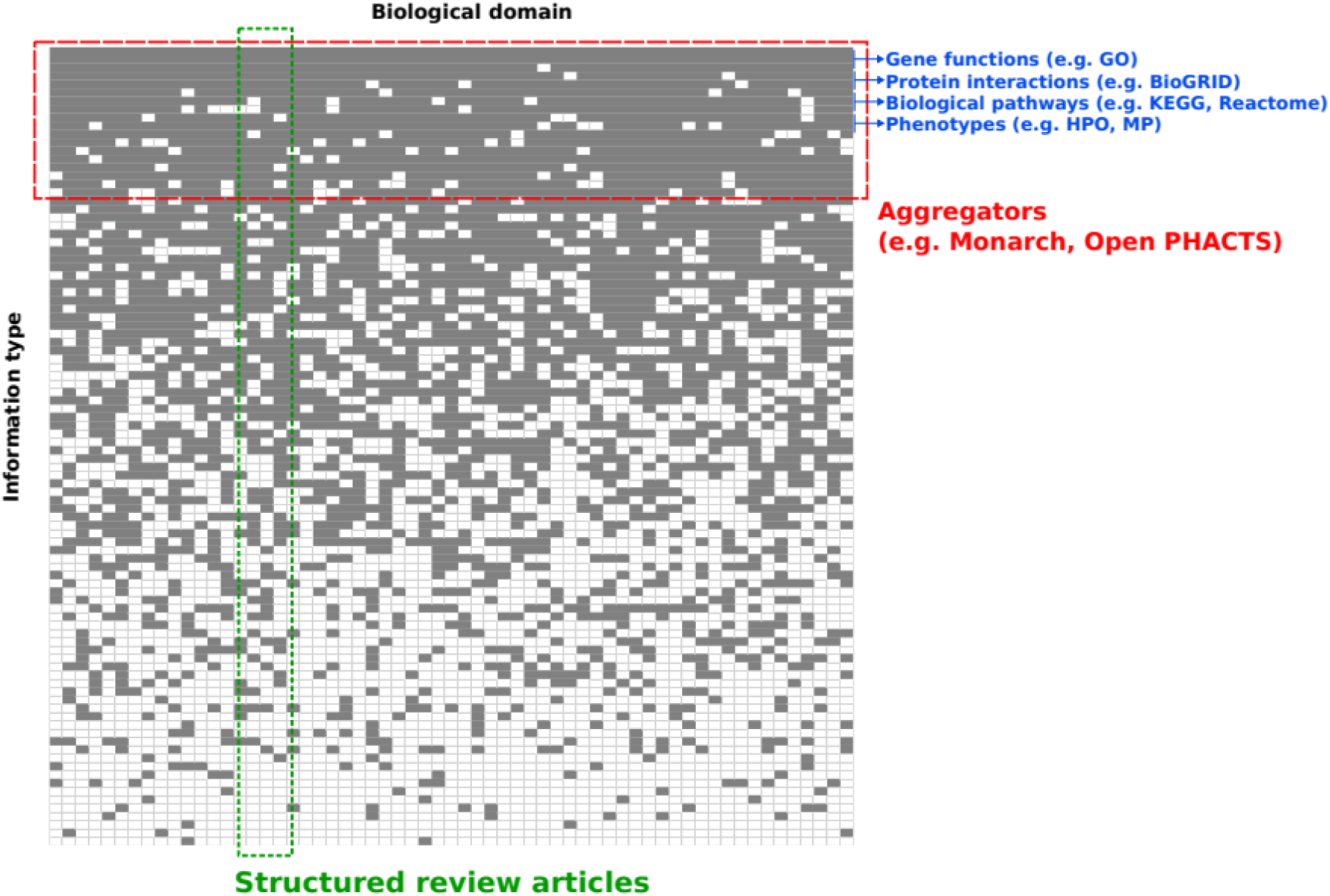
Conceptual overview of structured review articles. The landscape of biomedical information resources is heterogeneous and broad. Most data curation efforts focus on one (or a few) types of information, and on organizing that type of information across the entire literature without bias to the specific biological domain. This work focuses on the complementary challenge of organizing a diverse and heterogeneous graph of knowledge in a relatively defined domain area, and performing this integration in a way that is amenable to computational analysis. We call this type of effort a “structured review article”.

Knowledge graphs are computer-readable semantic representations of information, where concepts are encoded as nodes, and the relationships between those concepts are represented as edges. Knowledge graphs make it easy to integrate information from many sources, to explore heterogeneous information within a single data model, and to infer new relationships via efficient queries. Knowledge graphs have been used to organize background knowledge for data interpretation and hypothesis generation in a wide variety of contexts (6–9,4,10–12).

We propose structured review articles as knowledge graphs focused on specific disease and research questions. The goal of a structured review article is to organize relevant knowledge to make it interpretable and queryable by humans and computers (**Figure 1**). A structured review is similar to a systematic review in that it attempts to summarize current knowledge and evidence relevant to a research question, but it is different in that the knowledge is assembled in a modern and computable fashion. The benefit of this transformation is that the knowledge is computationally accessible and efficiently processable. This allows for the application of graph and artificial intelligence algorithms and tools, and better promotes the FAIR principles (13,14). As a proof-of-concept, we worked with researchers studying NGLY1 Deficiency (DOID:0060728), an ultra-rare disease first reported in 2012 that affects less than 100 patients worldwide with no treatment (15). They previously found evidence of a genetic association between NGLY1 and Aquaporins (16) where multiple aquaporins’ transcription was reduced in NGLY1 deficient cells by an unknown mechanism. Here, we explored the use of knowledge graphs as structured review articles to identify plausible regulatory mechanisms to explain these observations.

To demonstrate the value of this concept, we created a structured review article of NGLY1 Deficiency. In contrast to traditional free-text review articles, this structured review exists as a living knowledge graph that is curated by humans and accessible to computational analyses. Construction of this structured review was based on an iterative cycle of defining a research question, ingesting relevant data resources, and querying the resulting knowledge graph. Essential to this process was a close collaboration and iterative design with domain experts in NGLY1 Deficiency. To generalize this process to any domain area, we created tools to assist the creation and exploration of focused knowledge graphs. We also created a tool for community curation and contribution. We show how structured reviews are efficient knowledge structures for access and usability for humans and computers that may be particularly useful in rare disease research to springboard hypothesis building, to identify potential collaborations, and suggest potential testable hypotheses for drug discovery or repurposing.

## Methodology

### Biomedical data

To construct our NGLY1 Deficiency-focused knowledge graph, we utilized the following structured knowledge and data resources:

- We used Wikidata (http://wikidata.org/) to retrieve metadata for the biocurated network such as identifiers (IDs) from different vocabularies or entity cross-references, human readable labels, synonyms or descriptions. Wikidata is a project of the Wikimedia Foundation that enables the collaborative construction of a centralized graph database. Wikidata contains biomedical knowledge populated automatically from trusted authorities such as NCBI’s Entrez Gene, PubChem, and the Human Disease Ontology (17). Using the Wikidata SPARQL API (http://query.wikidata.org) we retrieved data from the 201703 version.
- We used the Monarch Initiative platform (3) to retrieve human and animal model biological data and metadata. The Monarch Initiative is developing a Knowledge Graph devoted to semantically integrate genomic, phenomic and related information from several species, tracking the evidence of the relationships. This integration is done with a clear emphasis to translate biomedical curated knowledge from animal model to human biology. Using the Monarch Biolink API (https://api.monarchinitiative.org/api/) we retrieved data from the 201901 version.
- We used the tftargets R package (https://github.com/slowkow/tftargets) and the Molecular Signature Database (MSigDb) (18,19) to retrieve human transcription factors (TFs) and their associated target genes data and metadata. tftargets aggregates experimental and curated gene regulatory information from the TRED (20), ENCODE (21), Neph2012 (22), and TRRUST (23) databases. We also retrieved regulatory relationships from MSigDB, a collection of annotated gene sets (the C3:TFT sub-collection v6.1) (24).
- We included one RNA-Seq dataset on a Drosophila model of NGLY1 Deficiency (25). To create lists of differentially expressed genes, we filtered for absolute fold change > 1.5 and false discovery rate (FDR) < 0.05.
- We used the BioThings MyGene.info API (http://mygene.info/) (26,27) to annotate synonyms, name, and description node attributes. We queried mygene.info services on 2019-01.
- We used Human Metabolome Database (HMDB) (28) to manually extract facts related to the GlcNAc metabolite. The Human Metabolome Database (HMDB) is a freely-available electronic database containing detailed chemical, clinical, and biological information about small molecule metabolites found in the human body. We used the version 2017-05.

### ID Normalization

To normalize entities and relations from different data models we used a variety of methods. For normalization of curated data to Monarch model we used the Wikidata SPARQL endpoint to retrieve cross-references to map entities. Diseases were linked to MONDO IDs by adding an extra ‘skos:exactMatch’ relationship. We used the MONDO ontology (http://obofoundry.org/ontology/mondo.html) to link disease IDs and retrieve node metadata. We used the OWL file version 2018-04-15 (http://purl.obolibrary.org/obo/mondo/releases/2018-04-15/mondo.owl). Genes were normalized by using BioThings MyGene.info API. We queried mygene.info services on 2019-01. Specifically, for normalization of curated data we queried for NCBI Gene IDs to HGNC and UniProt to HGNC, for normalization of regulatory genes we queried for Gene symbol to NCBI Gene IDs and to HGNC, and for NCBI Gene ID to HGNC and to Gene Symbol. The semantics of all relationships were manually mapped to the ontologies used in Monarch model.

### The BioKnowledge Reviewer Library

We created a library using the Python 3 programmatic language to reproduce the creation protocol of the structured review in a workflow. Functionality was guided by knowledge and reasoning of NGLY1 researchers. Under the research question, experts formulated a priori the specific questions they wanted to explore in the graph. The library is designed to give flexibility on the construction of the review network by choosing, concatenating and merging topic-specific networks. It allows to build reviews in a modular way by steps in the workflow and by different topics. We adopted a modular approach for their management to facilitate their reusability, update, and consistency checking. In **Figure 2** we illustrate the architecture of the system composed of four components. The edges component contains libraries with functions to collect, normalize, format the information and data we want to integrate. The graph component contains functions to integrate and create the knowledge graph. The Neo4j component contains the module to import the graph into Neo4j. Finally, the hypothesis-generation component contains the modules to query the graph, structure the resulting semantic paths, and extract summaries to analyse connections and the evidence. This programming library is versioned in GitHub to enable the community to add new functionality required to review new pieces of knowledge and apply it to answer more diverse types of questions. Availability: Library: https://github.com/SuLab/bioknowledge-reviewer Workflow notebook: https://github.com/SuLab/bioknowledge-reviewer/blob/master/bioknowledge_reviewer/graph_v3.2_v20190616.ipynb

**Figure 2.**
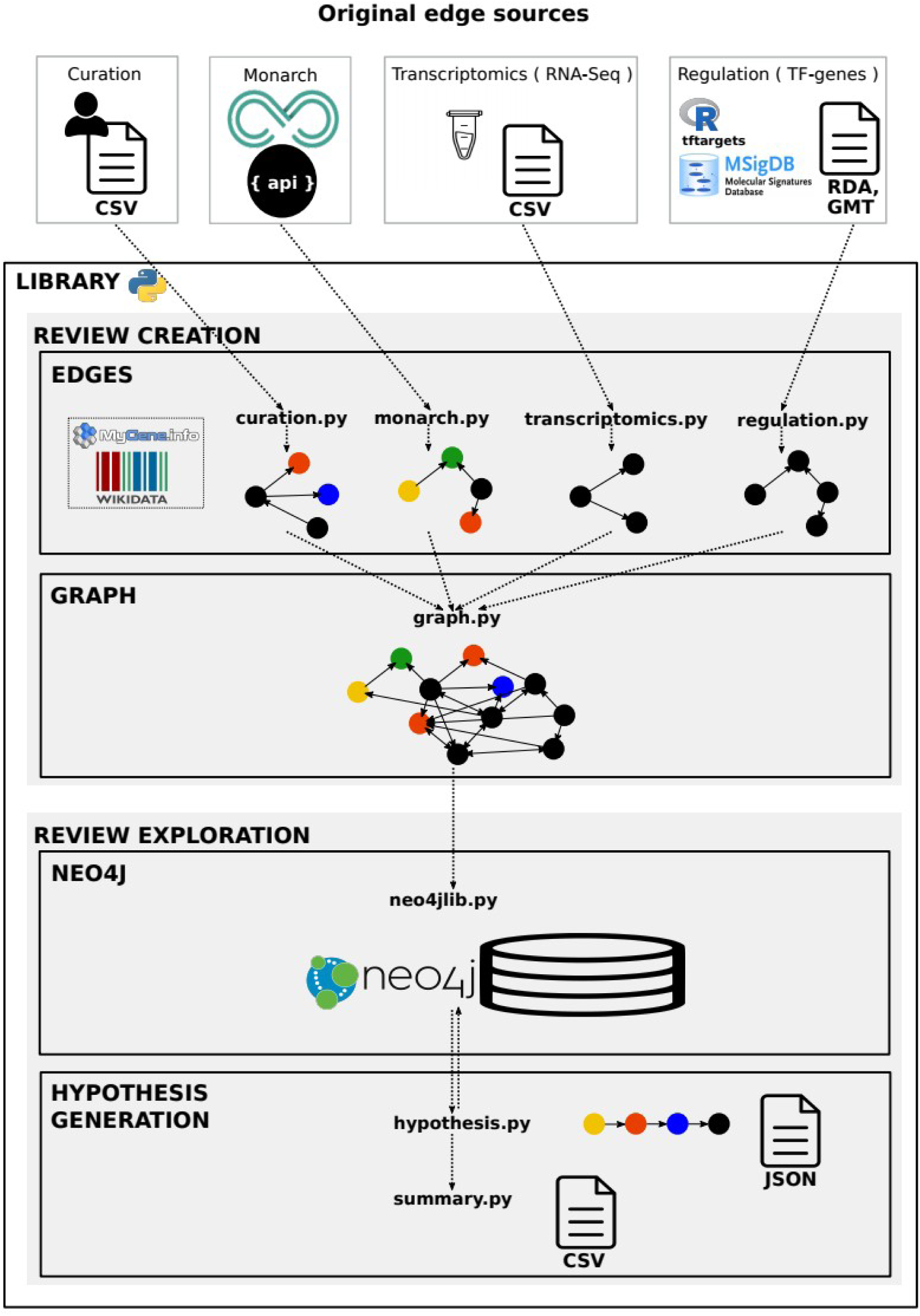
Library architecture. Architecture of the system based on four components.

### Data storage and mining

We used the Neo4j graph database framework for storage, management and mining of structured data. The graph database approach has been shown to facilitate management and exploration of biomedical knowledge (29). Neo4j enables users to query the graph using the Cypher query language, either through an API or a graphical user interface. All data were imported into the Neo4j Community Server v3.5.3.

### Evaluation

To evaluate the disease-based biocuration, we utilized the Semantic MEDLINE Database (SemMedDB) (30). SemMedDB is a repository of semantic relations sentence-based extracted from the titles and abstracts of all PubMed citations by a general knowledge-based text mining system called SemRep (31). We used version semmedVER31_R, which contains information about approximately 94.0 million relations from all of PubMed citations (about 27.9 million citations).

## Results

### 1 Construction of the knowledge graph

The overarching goal of our study was to generate mechanistic hypotheses for recent experimental observations. Specifically, we sought to explain the phenotypic effects of aquaporins on cellular phenotypes of NGLY1 Deficiency. Researchers found a transcriptional regulation link between NGLY1, ENGASE and AQP1 (16). However, the mechanism of this effect on a molecular level was not clear. Therefore, to identify plausible potential mechanisms to explain this observation and others like it, we iteratively constructed a knowledge graph that focused on information relevant to the NGLY1 gene, NGLY1 Deficiency, and aquaporins.

### Domain expert knowledge

We seeded our graph with nine key concepts as initial nodes – seven genes (NGLY1 human (HGNC:17646), AQP1 human (HGNC:633), AQP1 mouse (MGI:103201), ENGASE human (HGNC:24622), NFE2L1 human (HGNC:7781), AQP3 human (HGNC:636), AQP11 human (HGNC:19940)), one metabolite (GlcNAc (HMDB:HMDB00215)), and the disease NGLY1 Deficiency (OMIM:615273). NFE2L1 is a transcription factor recently discovered to be dependent on NGLY1 (32–34). Using these nodes as seeds, we then sought to retrieve as much biomedical information as possible on these entities. We first consulted the knowledge base created by the Monarch Initiative. We expanded our network to include to all concepts with an explicit relationship to one of our nine seed nodes. This expanded network included 713 nodes (including 234 genes, 80 diseases/phenotypes, 174 pathways, 111 tissues, 49 gene variants, 65 genotypes) and 6756 edges. Although the Monarch-derived network provided an important foundation for our work, we found that Monarch expansion alone did not represent all the knowledge we expected. Therefore, we then performed a targeted expansion of our network via three strategies.

### Biocuration

To include the most recent findings described in the literature around the molecular basis and clinical description of the disease, we curated two scientific papers that together compiled biomedical knowledge relevant to NGLY1 Deficiency. Together, the papers by Enns *et al*. in 2014 (35) and Lam *et al*. in 2017 (36) captured the known molecular biology involved in the disorder and the most recent and complete characteristics of the clinical phenotypic spectrum. Based on these two papers, we added to our knowledge network 101 phenotypes linked to NGLY1 Deficiency and an additional 142 biological relationships joining genes, variants and functions. This work also directly led to the creation of a term for NGLY1 Deficiency in the Human Disease Ontology (37), and the creation of 45 new phenotype terms in the Human Phenotype Ontology (38).

To evaluate the relevance of this curation effort, we compared our results with text mined statements in SemMedDB. Our manual curation effort resulted in more relationships (243 curated versus 11 text mined statements) and with more precise expressivity than via text mining. For instance, SemMedDB identified only four statements related to NGLY1 Deficiency phenotypes, and the information content was much less precise than our manual biocuration.

In addition, Owings *et al.* showed that GlcNAc is a metabolite with a potential key role in the molecular basis of NGLY1 Deficiency (25). As Monarch does not include metabolites in its knowledge base, we extracted edges related to GlcNAc from the Human Metabolome Database (28), KEGG (39), ChEBI (40), and ChEMBL (41). This work resulted in the addition of 362 edges and 302 nodes to our knowledge graph.

### Ortholog phenotypes

To increase the connectivity around NGLY1 and aquaporins, we included animal model information since the conserved biology could help explain the pathology in humans. We first added to our graph the orthologs for the seven seed genes, as well as the orthologs for all genes connected to any of the nine seed nodes. From these gene-ortholog edges, we then added all ortholog-phenotype relationships from Monarch. As a result we expanded our network with 246 new ortholog nodes, 570 new phenotype nodes, and 4,930 new edges.

### Transcription Factor regulation

To test the hypothesis that NGLY1 and aquaporins are mechanistically related through altered transcriptional regulation, we looked for data sources of known experimentally determined TF-gene relations. We also integrated a recently-published NGLY1 Deficiency fly model transcriptomic profile data set (25) into the review, and merged the collected TF-gene data. As a result we expanded our network with 9,723 TF-gene edges and 386 expression edges (including 4,226 new genes from which 640 are known TFs).

Finally, we again used the Monarch database to retrieve pairwise relationships between network nodes. The final knowledge graph contained 9,361 nodes, including 6,152 genes or proteins, 2,486 diseases or phenotypes, 355 pathways, 193 genotypes, 117 tissues, 50 variants, 7 chemical compounds and 1 organism, and contained 234,717 edges across 29 relationship types. Our final NGLY1-focused knowledge graph integrated data and knowledge derived from scientific literature, domain experts, databases and gene expression data.

### 1.1 Data model

The data model represents heterogeneous biomedical knowledge using common controlled vocabularies and ontologies in the Life Sciences to identify nodes and edges. Regarding node and edge human readable descriptions, we put special emphasis to be sufficient and efficient, i.e. the minimum but useful amount of information for biologists to understand the relational information, the entities involved and the supporting evidence backing each edge. For nodes we included an identifier (ID), a label, a name and a description, and for edges a property ID, a label, a property Uniform Resource Identifier (URI) to link to more detailed description of the semantics of the relation, a sentence supporting the relation extracted from the original scientific publication, and a reference URI to it to trace back the description of the relation in the original research study. A detailed description of the data model is available at ngly1graph.org.

## 2 Implementation and availability

### 2.1 Review creation by bioinformatic workflows

To create and analyse the structured review, we created a Python library and a jupyter notebook based on the knowledge graph construction protocol described above. The library allows any researcher to automate and reproduce the ingestion, integration of all data sources, and the creation of the knowledge graph in bioinformatic workflows. Every review can be created by mixing and remixing evolving knowledge on different topics, and it is versioned for use and reuse. Using the library, we can also derive structured hypotheses alongside each structured review, in the same workflow. This enables researchers: 1) to update the review iteratively avoiding redundant effort, and with own private datasets, 2) to reuse reviews to revisit the hypothesis with new findings, and 3) to share with other researchers for reproduction, discussion or curation or even as a citation. In addition to automating the exact work described in this manuscript, these programmatic tools also allow this general strategy to be generalized to other disease areas, and enable other developers to extend our work to new knowledge resources.

### 2.2 Data access

We stored the resulting graph in a Web-based application for dissemination, querying, visualization, and curation. This application is a hybrid between two technologies – Neo4j and Wikibase – with complementary strengths: knowledge navigation and knowledge contribution. Neo4j is a graph database with several useful features. First, Neo4j offers a powerful graph query language (“Cypher”) that enables any researcher to mine the database on our central server without having to set up any computer hardware or software of their own. Second, Neo4j also provides a Web-based graphical user interface for interactive database access and exploration. To provide simple starting points, we created Neo4j “guides” as basic tutorials, which contain templates for representative queries that can be extended and customized. And third, Neo4j utilizes a simple structured data format for both data import and export. This data structure facilitates the downstream reuse of the network using external tools and custom analyses.

The second component of our application is based on Wikibase, a system that enables living community curation and contribution of structured data. One of our key motivating design features was the ability for the NGLY1 community to contribute to a centralized community resource, but Neo4j does not have an easy mechanism for this purpose. Wikibase is the open source software that powers Wikidata, a crowdsourced effort to curate, manage, and share structured data. The Wikibase software offers manual editing via a Web-based interface, as well as automated editing via an API. Wikibase also includes detailed change tracking, and a SPARQL endpoint as the RDF query service a key technology for connecting with Linked Data and the Semantic Web.

Joining the Neo4j and Wikibase components in a single hybrid system is a continuously-running, real-time synchronization engine. This hybrid system combines the complementary strengths of each component – community contributions via Web-based and programmatic interfaces, a Web-based interface for graph visualization, and powerful query capabilities for discovery through two widely-used query languages. Finally, the structured review can be accessed via download of code and data, data as CSV or the interoperable RDF format.

## 3 Applications

### 3.1 Community curation

The NGLY1 community (and other rare disease communities in general) do not have the resources to sustain focused biocuration efforts. Therefore, we turned to community curation as a mechanism to continue the maintenance and expansion of the NGLY1 knowledge graph. Our interface for community curation is based on Wikibase, the software underlying Wikidata. This Wikibase extension allows anyone in the community to directly add and edit information to the NGLY1 knowledge graph through an online, graphical interface. Importantly, community curated data is fully structured at time of submission, making it fully integrated into the knowledge network.

This centralized, community-maintained resource will facilitate the exchange of knowledge across the entire NGLY1 community. Wikibase also makes these data interoperable with the broader ecosystem of Linked Data resources, which in turn will facilitate further reuse and additional data integration.

### 3.2 Hypothesis generation and exploration

A structured review brings together different topics that opens the possibility to unveil hypotheses by mining existing knowledge, as well as by offering additional ways to use existing knowledge to contextualize newly-generated data. Now the user can easily interrogate multi-dimensional and multi-domain knowledge and data in the structured review from a single endpoint. The user can traverse the NGLY1 Deficiency gene expression data in Drosophila, human TF-gene relationships, and the heterogeneous content on genes, pathways, phenotypes, or gene-gene interactions, and their cross-talk between different species to mine the gap of knowledge to generate hypotheses around the research question. Some example queries that explore potential mechanistic links between NGLY1 and AQP1 include:

1. Do NGLY1 and AQP1 (or related genes) share any phenotypes in knockout animals?
2. Do NGLY1 interacts with AQP1 transcriptional regulators?
3. Do AQP1 and phenotypes of NGLY1 Deficiency share related genes?

To illustrate a possible use of the NGLY1 Deficiency knowledge graph, we developed the research question of whether AQP1 gene may be important in the NGLY1 Deficiency phenotypes. To answer this question, biologists can interrogate the review as a graph with only two simple queries: a first query to find mechanistic links between NGLY1 and AQP1 gene expression, and a second query to find links between these mechanisms and the disease phenotypes. These queries are detailed in a Neo4j guide available at ngly1graph.org.

First, to identify mechanistic links between these two genes we formulated a query template based on the hypothesis that they are related via transcriptional regulation looking for an NGLY1 dependent transcription factor which regulates AQP1 or what we called herein the *regulatory hypothesis*. We found no direct regulatory links between the human NGLY1 gene and AQP1 gene. Therefore, we expanded our query to include regulatory links that were described in Drosophila orthologs (25) (**Figure 3A**).

**Figure 3.**
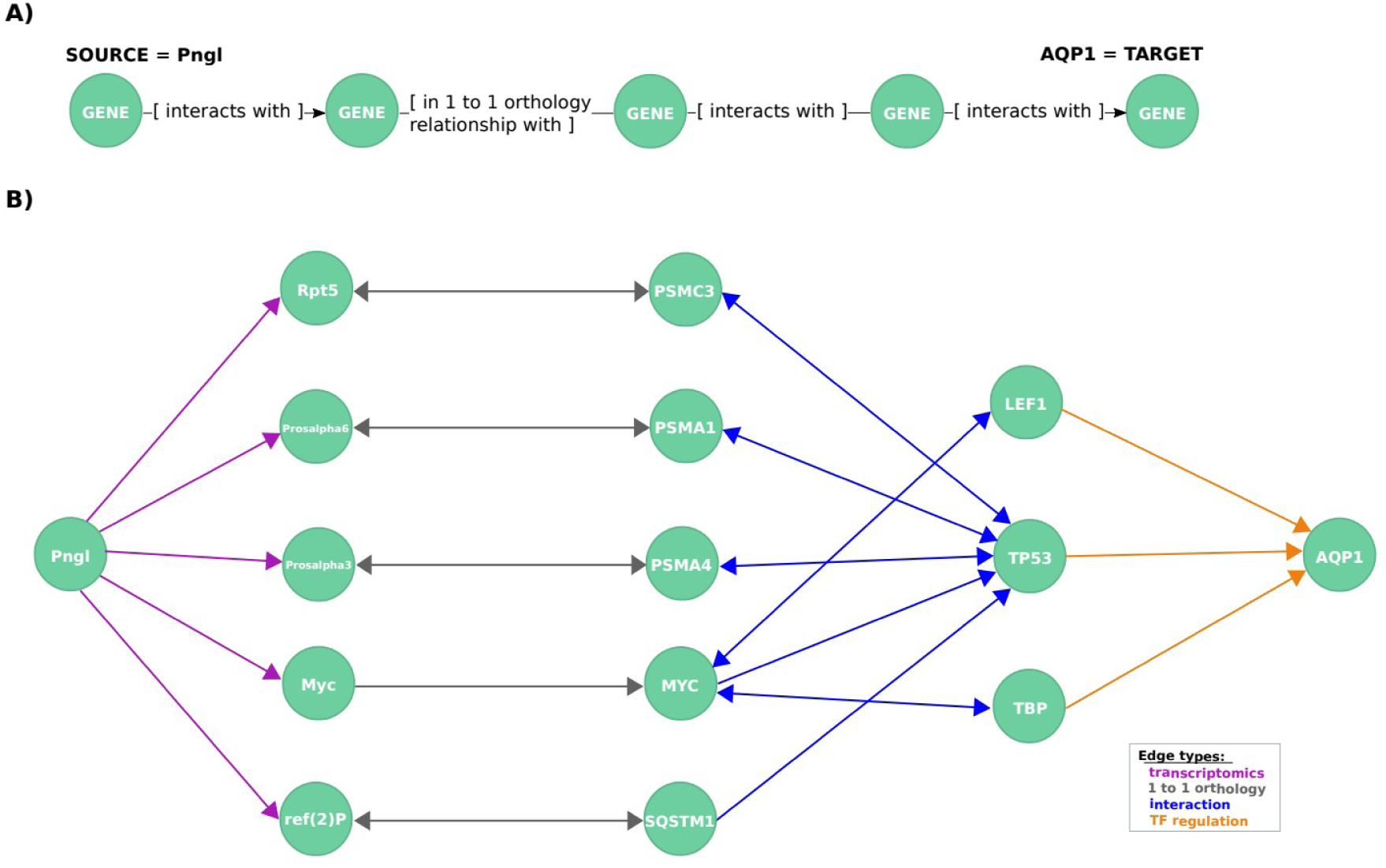
Exploration of mechanistic paths between NGLY1 and AQP1 based on the regulatory hypothesis. (A) First query topology for the regulatory hypothesis. We defined a path topology based on gene pathways of length four linking the NGLY1 ortholog in Drosophila (Pngl) with the human AQP1 gene. The bridging nodes and edges were based on transcriptional regulatory relationships in both Drosophila and human, plus orthology relationships between human and fly genes. (B) Mechanistic hypotheses resulted from the first query.

This query returned 19 paths, each of which represents a potential mechanistic hypothesis of how NGLY1 and AQP1 are related (**Figure 3B**). The genes TP53 (HGNC:11998), TBP (HGNC:11588) and LEF1 (HGNC:6551) are candidate regulators identified by the query. TP53 interacts via the proteasome complex (PSMA1 (HGNC:9530), PSMA4, PSMC3 (HGNC:9549)), SQSTM1 (HGNC:11280) a multifunctional protein that binds ubiquitin, and MYC (HGNC:7553), which is a phosphoprotein also related to TBP and LEF1. Further network exploration unveiled that TP53 interacts with TBP (42–46), and interestingly, that TBP is an interactor of MEF2A (HGNC:6993) (31), a known transcriptional regulator of AQP1 (24) and a member of a TF protein family recently associated with AQP1 transcription regulation (47).

To further explore this regulatory hypothesis, we queried the knowledge graph for relationships between phenotypes associated with NGLY1 Deficiency and the candidate genes identified in the previous query (**Figure 4A**). This query was designed to prioritize those candidate genes based on available prior evidence linking them to NGLY1 Deficiency phenotypes.

**Figure 4.**
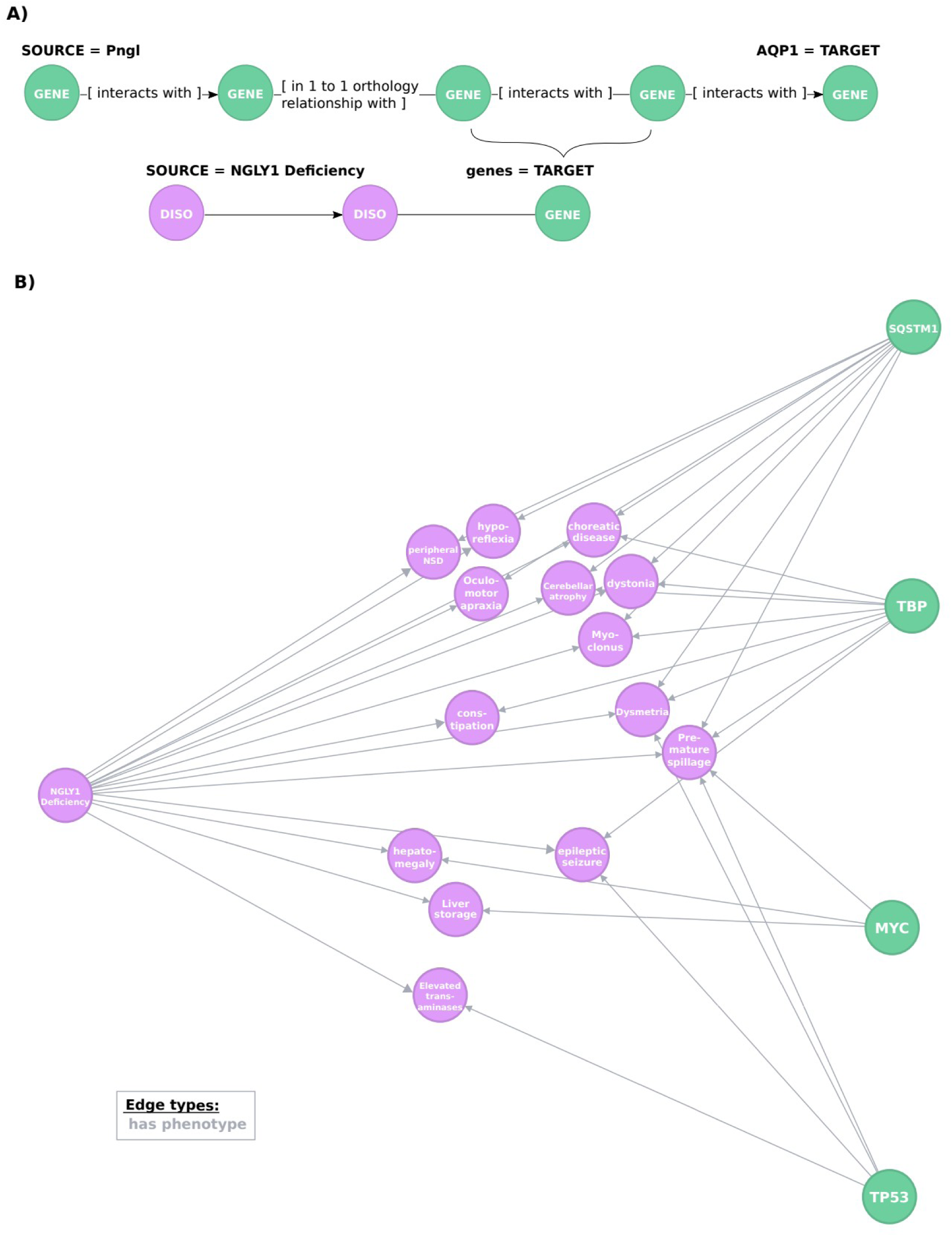
Exploration of the evidence relating candidate regulators of AQP1 to NGLY1 Deficiency phenotypes. (A) Second query topology for the AQP1 regulation-disease phenotypes shared genetic basis hypothesis. (B) Hypotheses resulted from the second query. All edges are of type ‘has phenotype’.

This query returned 30 paths that link AQP1 to 14 NGLY1 Deficiency phenotypes through genes involved in the regulatory hypothesis (**Figure 4B**). These paths highlighted a potential mechanistic role for four candidate genes: SQSTM1 (HGNC:11280), MYC (HGNC:7553), TP53 (HGNC:11998), TBP (HGNC:11588). For instance, SQSTM1 encodes a protein with regulatory activity on the inflammatory/immune responses related to the nuclear factor kappa-B signalling pathway, which is linked to cancer and nervous system processes such as synaptic plasticity and learning (48–51). The query results show that SQSTM1 has previously been linked to several phenotypes associated with NGLY1 Deficiency, including ‘Cerebellum atrophy’, ‘peripheral nervous system disease’, or interestingly ‘Dysmetria’, a phenotype caused by lesions in the cerebellum or proprioceptive nerves that lead to the cerebellum that coordinate visual, spatial and other sensory information with motor control (52). From the first query we can see that SQSTM1 interacts with TP53, which in turn it is also linked to ‘Dysmetria’.

In the Neo4j interface, users can interactively visualize any of the paths identified in these queries, check entity attributes such as a human readable description, and explore the evidence and context of each statement accessing the supporting reference through its URI. Users can also extract a table summarizing interesting path features like the most common genes from the transcriptome or the most common TFs (see an example in the Neo4j Guide).

The examples described here are just two representative queries that demonstrate the power of mining a semantically-precise knowledge graph. The Cypher query language offers powerful capabilities to harness the knowledge in our structured review. Users can formulate Cypher query templates corresponding to their biological question. Cypher templates can handle a broad spectrum of queries, from very precise queries that correspond to specific mechanistic hypotheses, to open-ended queries that flexibly retrieve paths that incorporate arbitrary types of connectivity. More example queries using the Cypher query language are provided in the advanced Neo4j guide available from ngly1graph.org.

To explore this research question without the NGLY1 graph, researchers would have to explore several databases and make time-consuming pre-processing steps and analysis on the results of these searches. Instead, the knowledge graph we have created enables researchers to query the graph with complex questions, to traverse different domains and databases in one single query, and to explore disease-specific hypotheses and evidence. These extracted NGLY1-AQP1 regulatory mechanistic links provide a knowledge foundation for bench researchers to create an informed regulatory hypothesis to evaluate in the laboratory.

## Discussion

To enable knowledge exploration and exploitation for researchers working on a specific question, we explored the use of knowledge graphs as structured review articles. With our approach we built the first review article for NGLY1 Deficiency research, and we demonstrated that is now an actionable knowledge resource for the whole community. This resource supports knowledge discovery and dissemination, and it facilitates collaboration between experimental researchers and bioinformaticians.

This work was motivated by the goal of identifying mechanistic hypotheses for an experimental observation. While this general procedure is not unique to our effort, we have incorporated two features that we believe make this work a notable contribution in this area. First, we have focused on creating a structured review article, which is distinct from other review articles in that the output is computationally accessible, and also distinct from other structured data integration efforts because of its relatively narrow and deep focus on a particular domain area. Second, we have published this structured review article to the community in the form of a centralized resource (accessible at http://ngly1graph.org/contribute/), a living knowledge graph that can be continually refined by community curation.

Typically structured knowledge resources are created by curation efforts. On one hand, there are homogeneous edge-specific silos such as GO, BioGRID, STRING, HPRD, UniProt, Reactome, KEGG, or MP (2,1,53–56,39,57). On the other hand, there are heterogeneous data integration knowledge bases such as the Monarch knowledge graph (3). Curated resources are vast but incomplete because the majority of the wealth of knowledge is unstructured since expert curation cannot keep the pace of scientific production.

To organize this knowledge, several text mined heterogeneous knowledge bases have been developed (30,58). Text mined resources are comprehensive but without the semantic specificity required to be useful for the rare disease field. Also heterogeneous knowledge bases typically have a broad scope to get a systems level understanding, but lack a focus on domain-specific knowledge to address a specific question.

A structured review article helps to mine the gap of knowledge where other resources are incomplete or not expressive enough for the domain or question to solve. Here, we created a framework to aid in the construction of a new knowledge resource to synthesize information focused on a specific research question. These tools facilitate the integration of data from diverse heterogeneous resources, from manual curation, to biomedical databases, to experimental data, to expert knowledge. This knowledge integration framework produces complete research objects, i.e. a workflow with the data, the code, the graph and structured hypotheses. These research objects promote more efficient research and reproducible science. And relative to traditional review articles, structured review articles are more Findable, Accessible, Interoperable, and Reusable (59,60).

This work focused on NGLY1 Deficiency, an ultra-rare disease that has been diagnosed in fewer than 100 children worldwide. But, the principles and tools developed herein are generalizable to other domain areas, and we believe they will be of particular interest and utility to the rare disease community. The community curation application enables living structured review and promotes the creation of FAIR content from the time of its inception. In areas where investments in data infrastructure are modest, these tools will facilitate synergy between experimental and computational biologists, and between data curators and data miners.

## Acknowledgements

The authors thank Dr Mitali Tambe and Professor Hudson H. Freeze from the Freeze Lab in the Human Genetics Program at Sanford-Burnham-Prebys Medical Discovery Institute for their input, collaboration and help. We acknowledge the NCATS Biomedical Data Translator Hackathon 2018 in La Jolla for the inspiration and input. The FAIRness of a prototype of the library was evaluated at the NBDC/DBCLS BioHackathon 2018 in Matsue. We thank Jiwen Xin, Sébastien Lelong and Chunlei Wu from the Wu Lab at Scripps Research for their help.

## References

1. Oughtred, R., Stark, C., Breitkreutz, B.-J., et al. (2019) The BioGRID interaction database: 2019 update. Nucleic Acids Res., 47, D529–D541.

2. Oughtred, R., (2019) The Gene Ontology Resource: 20 years and still GOing strong. Nucleic Acids Res., 47, D330–D338.

3. Mungall, C. J., McMurry, J. A., Köhler, S., et al. (2017) The Monarch Initiative: an integrative data and analytic platform connecting phenotypes to genotypes across species. Nucleic Acids Res., 45, D712–D722.

4. Jupp, S., Malone, J., Bolleman, J., et al. (2014) The EBI RDF platform: linked open data for the life sciences. Bioinformatics, 30, 1338–1339.

5. Ratnam, J., Zdrazil, B., Digles, D., et al. (2014) The Application of the Open Pharmacological Concepts Triple Store (Open PHACTS) to Support Drug Discovery Research. PLOS ONE, 9, e115460.

6. Leach, S. M., Tipney, H., Feng, W., et al. (2009) Biomedical Discovery Acceleration, with Applications to Craniofacial Development. PLOS Comput. Biol., 5, e1000215.

7. Cohen, T., Whitfield, G. K., Schvaneveldt, R. W., et al. (2010) EpiphaNet: An Interactive Tool to Support Biomedical Discoveries. J. Biomed. Discov. Collab., 5, 21–49.

8. Callahan, A., Dumontier, M. and Shah, N. H. (2011) HyQue: evaluating hypotheses using Semantic Web technologies. J. Biomed. Semant., 2, S3.

9. Liekens, A. M., De Knijf, J., Daelemans, W., et al. (2011) BioGraph: unsupervised biomedical knowledge discovery via automated hypothesis generation. Genome Biol., 12, R57.

10. Livingston, K. M., Bada, M., Baumgartner, W. A., et al. (2015) KaBOB: ontology-based semantic integration of biomedical databases. BMC Bioinformatics, 16, 126.

11. Messina, A., Fiannaca, A., La Paglia, L., et al. (2018) BioGraph: a web application and a graph database for querying and analyzing bioinformatics resources. BMC Syst. Biol., 12.

12. Kawashima, S., Katayama, T., Hatanaka, H., et al. (2018) NBDC RDF portal: a comprehensive repository for semantic data in life sciences. Database J. Biol. Databases Curation, 2018.

13. Elliott, J. H., Turner, T., Clavisi, O., et al. (2014) Living Systematic Reviews: An Emerging Opportunity to Narrow the Evidence-Practice Gap. PLOS Med., 11, e1001603.

14. Shokraneh, F. (2019) Reproducibility and replicability of systematic reviews. World J. Meta-Anal., 7, 66–71.

15. Need, A. C., Shashi, V., Hitomi, Y., et al. (2012) Clinical application of exome sequencing in undiagnosed genetic conditions. J. Med. Genet., 49, 353–361.

16. Tambe, M., Ng, B. G. and Freeze, H. H. (2019) N-Glycanase 1 Regulates Aquaporins Independent of Its Enzymatic Activity. N-Glycanase 1 Regulates Aquaporins Independent of Its Enzymatic Activity; SSRN Scholarly Paper ID 3411247; Social Science Research Network, Rochester, NY, (2019).

17. Burgstaller-Muehlbacher, S., Waagmeester, A., Mitraka, E., et al. (2016) Wikidata as a semantic framework for the Gene Wiki initiative. Database, 2016.

18. Subramanian, A., Tamayo, P., Mootha, V. K., et al. (2005) Gene set enrichment analysis: A knowledge-based approach for interpreting genome-wide expression profiles. Proc. Natl. Acad. Sci. U. S. A., 102, 15545–15550.

19. Liberzon, A., Birger, C., Thorvaldsdóttir, H., et al. (2015) The Molecular Signatures Database Hallmark Gene Set Collection. Cell Syst., 1, 417–425.

20. Jiang, C., Xuan, Z., Zhao, F., et al. (2007) TRED: a transcriptional regulatory element database, new entries and other development. Nucleic Acids Res., 35, D137–D140.

21. The ENCODE Project Consortium (2012) An integrated encyclopedia of DNA elements in the human genome. Nature, 489, 57–74.

22. Neph, S., Stergachis, A. B., Reynolds, A., et al. (2012) Circuitry and Dynamics of Human Transcription Factor Regulatory Networks. Cell, 150, 1274–1286.

23. Han, H., Shim, H., Shin, D., et al. (2015) TRRUST: a reference database of human transcriptional regulatory interactions. Sci. Rep., 5, 11432.

24. Xie, X., Lu, J., Kulbokas, E. J., et al. (2005) Systematic discovery of regulatory motifs in human promoters and 3′ UTRs by comparison of several mammals. Nature, 434, 338.

25. Owings, K. G., Lowry, J. B., Bi, Y., et al. (2018) Transcriptome and functional analysis in a Drosophila model of NGLY1 deficiency provides insight into therapeutic approaches. Hum. Mol. Genet., 27, 1055–1066.

26. Xin, J., Mark, A., Afrasiabi, C., et al. (2016) High-performance web services for querying gene and variant annotation. Genome Biol., 17, 91.

27. Wu, C., MacLeod, I. and Su, A. I. (2013) BioGPS and MyGene.info: organizing online, gene-centric information. Nucleic Acids Res., 41, D561–D565.

28. HMDB 4.0: the human metabolome database for 2018 | Nucleic Acids Research | Oxford Academic. HMDB 4.0: the human metabolome database for 2018 | Nucleic Acids Research | Oxford Academic https://academic.oup.com/nar/article/46/D1/D608/4616873 (accessed Jun 19, 2019).

29. Lysenko, A., Roznovat, I. A., Saqi, M., et al. (2016) Representing and querying disease networks using graph databases. BioData Min., 9, 23.

30. Kilicoglu, H., Shin, D., Fiszman, M., et al. (2012) SemMedDB: a PubMed-scale repository of biomedical semantic predications. Bioinformatics, 28, 3158–3160.

31. Rindflesch, T. C. and Fiszman, M. (2003) The interaction of domain knowledge and linguistic structure in natural language processing: interpreting hypernymic propositions in biomedical text. J. Biomed. Inform., 36, 462–477.

32. Lehrbach, N. J., Breen, P. C. and Ruvkun, G. (2019) Protein Sequence Editing of SKN-1A/Nrf1 by Peptide:N-Glycanase Controls Proteasome Gene Expression. Cell, 177, 737-750.e15.

33. Tomlin, F. M., Gerling-Driessen, U. I. M., Liu, Y.-C., et al. (2017) Inhibition of NGLY1 Inactivates the Transcription Factor Nrf1 and Potentiates Proteasome Inhibitor Cytotoxicity. ACS Cent. Sci., 3, 1143–1155.

34. Lehrbach, N. J. and Ruvkun, G. (2016) Proteasome dysfunction triggers activation of SKN-1A/Nrf1 by the aspartic protease DDI-1. eLife, 5, e17721.

35. Enns, G. M., Shashi, V., Bainbridge, M., et al. (2014) Mutations in *NGLY1*cause an inherited disorder of the endoplasmic reticulum–associated degradation pathway. Genet. Med., 16, 751–758.

36. Lam, C., Ferreira, C., Krasnewich, D., et al. (2017) Prospective phenotyping of NGLY1-CDDG, the first congenital disorder of deglycosylation. Genet. Med., 19, 160–168.

37. Kibbe, W. A., Arze, C., Felix, V., et al. (2015) Disease Ontology 2015 update: an expanded and updated database of human diseases for linking biomedical knowledge through disease data. Nucleic Acids Res., 43, D1071–D1078.

38. Expansion of the Human Phenotype Ontology (HPO) knowledge base and resources | Nucleic Acids Research | Oxford Academic. Expansion of the Human Phenotype Ontology (HPO) knowledge base and resources | Nucleic Acids Research | Oxford Academic https://academic.oup.com/nar/article/47/D1/D1018/5198478 (accessed Jun 19, 2019).

39. Kanehisa, M., Furumichi, M., Tanabe, M., et al. (2017) KEGG: new perspectives on genomes, pathways, diseases and drugs. Nucleic Acids Res., 45, D353–D361.

40. Hastings, J., Owen, G., Dekker, A., et al. (2016) ChEBI in 2016: Improved services and an expanding collection of metabolites. Nucleic Acids Res., 44, D1214–9.

41. Gaulton, A., Hersey, A., Nowotka, M., et al. (2017) The ChEMBL database in 2017. Nucleic Acids Res., 45, D945–D954.

42. Seto, E., Usheva, A., Zambetti, G. P., et al. (1992) Wild-type p53 binds to the TATA-binding protein and represses transcription. Proc. Natl. Acad. Sci., 89, 12028–12032.

43. Sommer, M. H., Scully, A. L. and Spector, D. H. (1994) Transactivation by the human cytomegalovirus IE2 86-kilodalton protein requires a domain that binds to both the TATA box-binding protein and the retinoblastoma protein. J. Virol., 68, 6223–6231.

44. Cvekl, A., Kashanchi, F., Brady, J. N., et al. (1999) Pax-6 interactions with TATA-box-binding protein and retinoblastoma protein. Invest. Ophthalmol. Vis. Sci., 40, 1343–1350.

45. Wu, Y., Mehew, J. W., Heckman, C. A., et al. (2001) Negative regulation of bcl-2 expression by p53 in hematopoietic cells. Oncogene, 20, 240.

46. Qadri, I., Iwahashi, M. and Simon, F. (2002) Hepatitis C virus NS5A protein binds TBP and p53, inhibiting their DNA binding and p53 interactions with TBP and ERCC3. Biochim. Biophys. Acta BBA - Mol. Cell Res., 1592, 193–204.

47. Jiang, Y., Liu, H., Liu, W., et al. (2016) Endothelial Aquaporin-1 (AQP1) Expression Is Regulated by Transcription Factor Mef2c. Moleucles Cells, 39, 292–298.

48. Albensi, B. C. and Mattson, M. P. (2000) Evidence for the involvement of TNF and NF-κB in hippocampal synaptic plasticity. B in hippocampal synaptic plasticity. Synapse, 35, 151–159.

49. Meffert, M. K., Chang, J. M., Wiltgen, B. J., et al. (2003) NF-κB in hippocampal synaptic plasticity. B functions in synaptic signaling and behavior. Nat. Neurosci., 6, 1072.

50. Vlahopoulos, S. A., Cen, O., Hengen, N., et al. (2015) Dynamic aberrant NF-κB in hippocampal synaptic plasticity. B spurs tumorigenesis: A new model encompassing the microenvironment. Cytokine Growth Factor Rev., 26, 389–403.

51. Vlahopoulos, S. A. (2017) Aberrant control of NF-κB in hippocampal synaptic plasticity. B in cancer permits transcriptional and phenotypic plasticity, to curtail dependence on host tissue: molecular mode. Cancer Biol. Med., 14, 254-270–270.

52. Spatial Attention Deficits in Patients with Acquired or Developmental Cerebellar Abnormality | Journal of Neuroscience. Spatial Attention Deficits in Patients with Acquired or Developmental Cerebellar Abnormality | Journal of Neuroscience http://www.jneurosci.org/content/19/13/5632.long (accessed Jun 19, 2019).

53. Szklarczyk, D., Gable, A. L., Lyon, D., et al. (2019) STRING v11: protein–protein association networks with increased coverage, supporting functional discovery in genome-wide experimental datasets. Nucleic Acids Res., 47, D607–D613.

54. Keshava Prasad, T. S., Goel, R., Kandasamy, K., et al. (2009) Human Protein Reference Database—2009 update. Nucleic Acids Res., 37, D767–D772.

55. Keshava Prasad, T. S.,(2019) UniProt: a worldwide hub of protein knowledge. Nucleic Acids Res., 47, D506–D515.

56. Fabregat, A., Jupe, S., Matthews, L., et al. (2018) The Reactome Pathway Knowledgebase. Nucleic Acids Res., 46, D649–D655.

57. Smith, C. L. and Eppig, J. T. (2009) The mammalian phenotype ontology: enabling robust annotation and comparative analysis. Wiley Interdiscip. Rev. Syst. Biol. Med., 1, 390–399.

58. Hristovski, D., Dinevski, D., Kastrin, A., et al. (2015) Biomedical question answering using semantic relations. BMC Bioinformatics, 16, 6.

59. Wilkinson, M. D., Dumontier, M., Aalbersberg, Ij. J., et al. (2016) The FAIR Guiding Principles for scientific data management and stewardship. Sci. Data, 3, 160018.

60. Brandizi, M., Singh, A., Rawlings, C., et al. (2018) Towards FAIRer Biological Knowledge Networks Using a Hybrid Linked Data and Graph Database Approach. J. Integr. Bioinforma., 15.

